# Trait-based aerial dispersal of arbuscular mycorrhizal fungi

**DOI:** 10.1101/2020.02.12.943878

**Authors:** V. Bala Chaudhary, Sarah Nolimal, Moisés A. Sosa-Hernández, Cameron Egan, Jude Kastens

## Abstract

- Dispersal is a key process driving local-scale community assembly and global-scale biogeography of plant symbiotic arbuscular mycorrhizal (AM) fungal communities. A trait-based approach could improve predictions regarding how AM fungal aerial dispersal varies by species.
- We conducted month-long collections of aerial AM fungi for 12 consecutive months in an urban mesic environment at heights of 20 m. We measured functional traits of all collected spores and assessed aerial AM fungal community structure both morphologically and with high-throughput sequencing.
- Large numbers of AM fungal spores were present in the air over the course of one year and these spores were more likely to exhibit traits that facilitate dispersal. Aerial spores were smaller than average for Glomeromycotinan fungi. Trait-based predictions indicate that nearly 1/3 of described species from diverse genera demonstrate the potential for aerial dispersal. Diversity of aerial AM fungi was relatively high (20 spore species and 17 virtual taxa) and both spore abundance and community structure shifted temporally.
- The prevalence of aerial dispersal in arbuscular mycorrhizas is perhaps greater than previously indicated and a hypothesized model of AM fungal dispersal mechanisms is presented. Anthropogenic soil impacts may initiate the dispersal of disturbance-tolerating AM fungal species and facilitate community homogenization.

## INTRODUCTION

Dispersal is a fundamental process influencing both large-scale biogeographical patterns and local community assembly. Plants in all terrestrial biomes on Earth form belowground symbioses with diverse communities of arbuscular mycorrhizal (AM) fungi, exchanging fixed carbon for improved access to soil resources (Smith & Read, 2008). These symbioses affect community, ecosystem, and global-scale patterns and processes (Johnson *et al.*, 2006) with function varying as AM fungal communities differentiate. Deterministic (niche-based) processes are important determinants of AM fungal community structure, but stochastic processes like dispersal can also influence AM fungal community structure and biogeographic patterns (Chaudhary *et al.*, 2008; Nielsen *et al.*, 2016). Knowledge regarding AM fungal dispersal mechanisms aids in the further incorporation of filamentous fungi into classic movement ecology models (Bielčik *et al.*, 2019). Furthermore, an improved understanding of AM fungal dispersal, and how it may vary among species, improves our ability to manage mycorrhizal symbioses in both natural and managed ecosystems (Hart *et al.*, 2018). For example, efforts to restore native mycorrhizal populations in anthropogenically disturbed soils could theoretically focus on species with limited dispersal capabilities, but such efforts require species-specific data on AM fungal dispersal to inform predictions.

Trait-based approaches are increasingly being utilized in ecology to shift from descriptive to predictive work (Messier *et al.*, 2010). Spores, the primary reproductive propagule for AM fungi, differ among species with respect to a suite of quantifiable morphological traits (e.g. intrinsic properties) that likely influence movement and dispersal capabilities. Arbuscular mycorrhizas notoriously form the largest single-cell fungal spores on Earth, with some species measuring larger than 1 mm in diameter and visible to the naked eye (Nicolson & Schenck, 1979). However, different species form spores up to two orders of magnitude smaller and, for a comparatively species-poor group, interspecific variation in AM fungal spore size is considerable (Aguilar-Trigueros *et al.*, 2018). Because spore size, to an extent, can be proportional to aerial dispersal predictors such as settling velocity (Kauserud *et al.*, 2008; Norros *et al.*, 2014), it could be a useful trait for making predictions regarding AM fungal dispersal. Other spore traits, such as surface ornamentation or color may also influence AM fungal aerial dispersal; species-specific pits or projections in spore surfaces could affect fluid drag (Roper *et al.*, 2008) and differences in the degree of spore melanization could be linked to stress tolerances such as UV radiation during aerial movement (Deveautour *et al.*, 2019). Patterns in fungal traits observed in aerially dispersed AM fungi have the potential to bring increased insight into predictions regarding which species or groups of species are more likely to disperse by wind or long distances.

The arbuscular mycorrhizal symbiosis occurs almost entirely belowground; AM fungi rarely form aboveground structures and the mechanisms of aerial dispersal are poorly understood. Studies have eluded to AM fungal aerial dispersal capabilities (Lekberg *et al.*, 2011; Davison *et al.*, 2015; Nielsen *et al.*, 2016), but since 1982 only four studies have documented the presence of AM fungal propagules in aerial samples through direct measurements. Propagules of AM fungi include asexual spores, hyphal fragments, and colonized root segments (Tommerup & Abbott, 1981; Biermann & Linderman, 1983). In laboratory tests using dry elutriation, the process of separating particles by size using an upward air current, AM fungal propagules and spores demonstrated the potential to remain suspended and travel aerially large distances (Tommerup, 1982). Large numbers of AM fungal spores were reported to be present in the air, able to move distances up to 2 km, and disperse in a non-random fashion associated with complex regional wind patterns (Warner *et al.*, 1987; Allen *et al.*, 1989). In a cross-biome study in North America, low numbers of AM fungal spores (10 min sampling period) were documented in the air predominantly in arid ecosystems (Egan *et al.*, 2014). These studies sampled air for periods of 10 min to 1 wk and uncovered very low fungal morphological diversity.

Still, significant knowledge gaps remain regarding our understanding of AM fungal dispersal (Bueno & Moora, 2019). Because the overwhelming majority of AM fungal species are hypogeous (but see Daniels & Trappe, 1979), their method for escaping the rhizosphere and becoming airborne to enable dispersal at multiple spatial scales is unclear. Also, it is unknown whether all AM fungal species disperse in a similar manner. This is important because interspecies variation in dispersal capabilities could contribute to differences in AM fungal community structure at local scales and biogeographic patterns at continent to global scales (Davison *et al.*, 2015). Furthermore, comparatively little is known about temporal patterns in AM fungal aerial dispersal. Temporal variation in dispersal is predicted because environmental factors that could influence aerial fungal dispersal, such as wind speed, temperature, and precipitation, also vary seasonally (Li & Kendrick, 1995). Temporal variation in aerial AM fungal dispersal is also predicted because of seasonal sporulation patterns that differ across species (Pringle & Bever, 2002). Temporal shifts in AM fungal sporulation are also linked to plant host phenology and other soil and habitat-related environmental variables (Liu *et al.*, 2009). Seasonal patterns in AM fungal sporulation, coupled with temporal fluctuation in meteorological conditions that impact dispersal, could result in a complex intersection of conditions that lead to stochastic AM fungal community assembly. The mechanisms of AM fungal dispersal are also important to understand from the applied perspective of restoring soil and ecosystem function. Commonly, in anthropogenically disturbed soils or primary succession situations, soils initially lack fungal propagules, but AM fungal communities passively develop over time (Johnson & McGraw, 1988; de León *et al.*, 2016; Chaudhary *et al.*, 2019). Less is known regarding which AM fungal species are more likely to disperse aerially or be first colonizers into disturbed soils (Hart *et al.*, 2018). Trait-based, predictive information regarding AM fungal dispersal could improve our understanding of the stochastic mechanisms that drive AM fungal biogeographical patterns and community assembly, as well as our ability to effectively manage mycorrhizas in ecosystems.

In this study, to examine trait-based AM fungal aerial dispersal, we assessed aerial AM fungal spore abundance, spore functional traits, and aerial AM fungal community structure (via both morphological and sequence-based methods) by collecting aerial samples from a mesic urban environment over the course of one year. Little is known about AM symbioses in urban ecosystems, including how dispersal and passive reestablishment of propagules into degraded urban areas might shape AM fungal community structure; because the outcome of AM symbioses can vary as fungal communities shift, studying AM fungal dispersal in cities has implications for efforts to improve urban sustainability (Chaudhary *et al.*, 2019). We also compare the measured traits of aerial spores to known traits for all described AM fungi present in the FUN^FUN^ fungal functional trait database (Zanne *et al.*, 2019). We predict that aerial AM fungal spores will possess traits more conducive to wind dispersal such as a smaller diameter than the average for all AM fungal species (Hypothesis 1). Smaller fungal spores are purported to travel further, at greater heights, and remain suspended for longer time periods (Wilkinson *et al.*, 2012; Norros *et al.*, 2014). We also predict that spore abundance will vary temporally over the course of the year and that aerial AM fungal community structure will differ seasonally (Hypothesis 2) because of divergent phenological strategies of different AM fungal species and their hosts (Gemma *et al.*, 1989; Pringle & Bever, 2002) and temporal variation in meteorological factors (e.g. wind speed, temperature). Finally, we predict low overall AM fungal diversity in aerial samples regardless of differences in observed aerial species depending on the detection method (spore morphology versus sequence-based) due to different types of AM fungal propagules (e.g. spores, hyphae) present in aerial samples (Hypothesis 3).

## MATERIALS AND METHODS

### Study Design

To examine trait-based AM fungal aerial dispersal, we conducted passive, isokinetic air sampling in the field for 12 consecutive months. Multiple BSNE dust collectors (Figure S1) were placed on a 20 m rooftop in a highly urbanized (population density ∼8,000/km^2^) location in Chicago, USA (41.88°N, 87.63°W). Named after the first successful prototype “Big Springs Number Eight”, BSNE dust collectors require no external power, can withstand extreme conditions, and isokinetically maintain sampling efficiency (Fryrear, 1986). A rotating weather vane keeps the collector oriented windward, and as air flows into the collection chamber, velocity slows, causing particles to drop into a collecting pan. We affixed metal covers to each chamber to keep collected material dry and prevent rain from entering chambers through the air outflow mesh. Previous field collections of aerial AM fungi have primarily used active filtration samplers that collect air samples for 2 hours; BSNE samplers allow for sampling periods of weeks or even months and have demonstrated success in aerobiological research (Johnson *et al.*, 2019). In our study, samples were collected after a period of 30 days. The direction of prevailing winds at the study site fluctuates seasonally with west/northwest winds in the winter, northeast winds in the spring, southwest/south winds in the summer and fall (IL Climate). The local climate is characterized as mesic temperate with cold windy winters and hot, humid summers (Hayhoe *et al.*, 2010).

BSNE samplers were deployed in the field in January 2017 and contents were collected monthly for laboratory analyses for 12 consecutive months. After the contents of each chamber were collected, chambers were thoroughly cleaned with ethanol prior to redeployment. Initially 3 BSNE collectors were deployed, but after 4 months, to boost replication, an additional 3 dust collectors were added to the study such that for January-April n=3, and for May-December n=6 resulting in 60 total observations ((4 months x 3 chambers) + (8 months x 6 chambers)). To explore meteorological predictors for aerial AM fungal dispersal, data on monthly maximum wind speed, average temperature, total precipitation, and maximum soil temperature for the year of 2017 were obtained from the Illinois Climate Network Water and Atmospheric Resources Monitoring Program (2015).

To explore potential plant phenological drivers of aerial AM fungal dispersal, we estimated end-of-season (EOS) week using a 2017 time series of normalized difference vegetation index (NDVI) data from the Advanced Very High resolution Radiometer (AVHRR) (Eidenshink, 1992; Eidenshink, 2006; United States Geologic Survey, 2019). These raster remote sensing data have 1 km spatial resolution and weekly temporal resolution, whereby each weekly scene is a maximum value composite of near-daily NDVI values obtained during the previous two weeks to minimize cloud impacts. For each pixel and each year, EOS was determined using the midpoint of the methods described in Zhang *et al.* (2003) and Yu *et al.* (2004) following time series preparation methods adapted from Wardlow et al. (2006).

### Spore and trait data collection

During monthly collection events, chambers were brought into the lab and collected material was carefully washed out of each chamber into a 25 µm sieve using distilled water. The contents of the sieve were washed and collected onto a 0.45 µm pore size gridded membrane filter using suction filtration. A subset of collected spores was gently scraped from the filter paper using fine forceps and permanently mounted onto microscope slides in poly-vinyl-lacto-glycerol (McKenney & Lindsey, 1987). Using a compound microscope (200-1000x magnification), spores were counted and identified to species using morphological characteristics from original species descriptions as well as more recent phylogenetic reclassifications (Redecker *et al.*, 2013). Assessment of AM fungal communities can be conducted using DNA-based techniques or through morphological identification of asexual spores; both approaches are informative and both have disadvantages (Öpik & Davison, 2016). One advantage of using spore morphological techniques is the ability to simultaneously collect information regarding functional traits of morphological species that can be used to inform predictions regarding the dispersal ecology of AM fungi. As such, we assessed aerial AM fungal spores microscopically for the physical traits of diameter and degree of melanization.

Diameter of each spore was directly measured using a ruled reticle, and degree of spore melanization was assigned an ordinal 1-5 rank, where 1 represents non-melanized, hyaline spores and 5 represents highly-melanized spores. Spores were also noted for either the presence or absence of surface ornamentation. Spore abundance reflects the total number of spores collected per sampling unit over the course of one month. Morphological species richness reflects the number of unique morphological species observed per sampling unit over the course of one month. A trait matrix was constructed for every spore encountered (47,162 observational rows by 3 trait columns); to examine patterns in spore traits over time, weighted averages were calculated to account for differences in abundances among morphological species.

### DNA-based assessment of AM fungal taxa

After a subset from each sample was removed for above spore analyses, samples were stored frozen (−20° C) on sterile membrane filters. To extract genomic DNA, membrane filters containing dust samples were cut into 4 mm strips using sterile scissors and placed into sample collection tubes for DNA extraction using the DNeasy PowerSoil Kit (QIAGEN, USA) according to standard protocol. Sequencing libraries were then prepared by the Environmental Sample Preparation and Sequencing Facility at Argonne National Laboratory (https://www.anl.gov/bio/environmental-sample-preparation-and-sequencing-facility), using the protocol as laid out in Morgan and Egerton-Warburton (2017). In brief, a segment of Glomeromycotinan small subunit ribosomal RNA (SSU rRNA) was amplified using the NS31 and AML2 primers (Simon *et al.*, 1992; Lee *et al.*, 2008) in a single-step PCR reaction. Both primers were modified for Illumina sequencing platforms by adding adapter sequences for annealing to Illumina flow cells, a “pad” sequence, and a two-base linker sequence, following the protocol of Caporaso *et al*. (2011).

Amplicons were then quantified using PicoGreen (Invitrogen) and a plate reader (Infinite 200 PRO, Tecan). Once quantified, volumes of each of the products were pooled into a single tube so that each amplicon is represented in equimolar amounts. This pool was then cleaned up using AMPure XP Beads (Beckman Coulter), and then quantified using a fluorometer (Qubit, Invitrogen). After quantification, the molarity of the pool was determined and diluted down to 2 nM, denatured, and diluted to a final concentration of 6.75 pM with a 10% PhiX spike for sequencing on the Illumina MiSeq. Amplicons were sequenced on a 301 bp x 12 bp x 301 bp MiSeq sequencer (Illumina Inc., San Diego, CA, USA) using custom sequencing primers. Post-sequencing, sequences were demultiplexed using Illumina’s bcl2fastq2 software.

### Bioinformatical analyses

From the sequencing run, a total of 1,020,046 sequences were generated from aerial samples. In addition to these sequences from aerial samples, we included 5,137,832 sequences during bioinformatic processing from two additional experiments that amplified AM fungal DNA from soil. These additional sequences were included because 1) amplicon libraries were generated using the same protocols as the present study, 2) they were sequenced on the same run as aerial samples, and 3) concurrent processing enabled a more robust estimation of sequencing error rates and more precise and reliable inference of amplicon sequence variants (ASVs). Sequences were first assembled into ASVs (Callahan *et al.*, 2017) with the DADA2 R package (Callahan *et al.*, 2016) using the standard pipeline (https://benjjneb.github.io/dada2/tutorial_1_8.html). In short, after primer removal, we trimmed sequences at 290 bp due to base-call quality degradation on the 3’ end of reads. The selected sequence trimming allowed for the minimal required overlap of 20 bp for the assembly of forward and reverse reads, given the expected amplicon length of approximately 560 bp (Öpik *et al.*, 2013). We then removed sequences with ambiguous bases, using maximum expected error thresholds of 2 and 5 for forward and reverse reads, respectively (Roy et al. 2019). ASVs were inferred on a global basis and sequences were merged. After chimera removal, we obtained 73,239 sequences assigned to 172 different ASVs.

The majority of aerial sequences (86%) were not successfully merged. Usually, this signifies issues with either the bioinformatics pipeline or the quality of sequences generated from the run. However, the same pattern was not observed in the sequences that were included from the other experiments where roughly 11% of sequences did not merge. Because we used identical primers and protocols to generate amplicon libraries for both the present study and the other experiments, we attribute this issue to the nature of our aerial samples. Specifically, because aerial samples likely harbor scant amounts of AM fungal DNA, we suspected the lack of merging resulted from a relative abundance of non-Glomeromycotinan DNA in aerial samples. As such, our primers may have often not found a target sequence, leading to non-specific amplification and ultimately resulting in sequences that could not be merged. A closer inspection of non-merged aerial forward sequences revealed that roughly 93% of these sequences, representing 96% of the forward non-merged reads, matched non-Glomeromycotina entries in the GenBank nucleotide database (https://www.ncbi.nlm.nih.gov/genbank/). Of these non-merged forward sequences, roughly 75% matched to Arachnida (85% of non-merged reads) and the rest to various fungal and metazoan sequences, confirming a high prevalence of non-specific amplification in our aerial samples. Likewise, 91% of non-matched reverse reads matched non-Glomeromycotina entries, representing 97% of the reverse non-merged reads. Here, again, 62% of the reverse non-merged sequences matched to Arachnida (87% of non-merged reads).

We then assigned ASVs using a BLAST+ search (Camacho *et al.*, 2009) against the MaarjAM database (Öpik *et al.*, 2010) accessed June 2019. Our required criteria for a match was defined as: sequence similarity ≥ 97%; alignment length not differing from the length of the shorter of our sequence and reference database sequences by > 5%; and a Blast e-value < 1e-50 following (Rodríguez-Echeverría *et al.*, 2017). Using these criteria, we assigned 54 (32%) of our ASVs to virtual taxa (VT) in MaarjAM. Subsequently, a second round of taxonomic identification was conducted by taking sequences that were not identified using the MaarjAM database and using a BLAST+ search against the GenBank nucleotide database. Criteria for a match was defined as: ≥90% sequence similarity, Blast e-value < 1e-50 and alignment length differing by < 10 nucleotides between the query and the subject. All sequences matched to either non-Glomeromycotina fungi, Metazoa, or Viridiplantae, further indicating either a high abundance of non-Glomeromycotina DNA within our aerial traps and/or a high prevalence non-specific amplification during amplicon library preparation.

### Data Analysis

Variation in traits for all observed aerial AM fungal spores was visualized using histograms and heatmaps. To test for statistical differences between location and shape of cumulative distributions of spore diameter of observed aerial spores and spore diameter for all described Glomeromycotina in the FUN^FUN^ database, we used Kolmogorov-Smirnov tests (Sokal & Rohlf, 1995). To test for differences in all AM fungal response variables across months, we conducted linear mixed effect models with month as the fixed model component and chamber identity as the random component (Zuur *et al.*, 2009). The random component was included in all models to account for non-independence and test for possible effects of variation in spore collection ability of the different chambers and chamber locations. Spore community structure was assessed with non-metric multidimensional scaling (NMS) using the Bray-Curtis distance measure and an iterative stepwise process to evaluate the dimensionality of the least stressful solution (McCune *et al.*, 2002). Differences in spore community structure among *a priori* seasonal groups (winter, spring, summer, fall) were assessed using single factor multi-response permutation procedure with 999 permutations (MRPP). Mixed effect models were conducted using the nlme package (Pinheiro *et al.*, 2014) and multivariate community analyses were conducted using the vegan package (Oksanen *et al.*, 2019). Pairwise comparisons among groups with a Bonferroni p-value adjustment were conducted using the RVAideMemoire package in R (Hervé, 2017).

Regarding molecular data, ASVs represented by only one read were kept only if i) they were also present in the bigger dataset and ii) they were not the only read assigned to their corresponding VT. In this way we kept 48 ASVs for further analysis. We aggregated samples by season and transformed sequence data into a presence/absence matrix. For the graphical representation of these data we used the package “ComplexHeatmap” (Gu *et al.*, 2016). We treated differences in numbers of reads per season as a biological signal and evaluated diversity and seasonality without any rarefaction method. For the validation of our conclusions we repeated our analyses on data rarefied to the lowest number of reads in a season (172, summer), by random sampling without replacement. Rarefaction, species accumulation curves and rarefaction curves were calculated in vegan (Oksanen et al., 2016). All analyses were conducted using R version 3.5.2 (R Core Team, 2016).

## RESULTS

### Aerial AM fungal traits

Over the course of one year, 47,162 total AM fungal spores were captured in aerial samplers (mean = 786/month ±782.9sd) at a height of 20 m in a highly urbanized, mesic environment. Aerial AM fungal spore traits varied substantially, with sizes ranging from 25 µm to 400 µm in diameter, melanization ranging from colorless to black, and the presence of both smooth and ornamented surfaces (Figure 1A, 2A). However, the overwhelming majority of aerial AM fungal spores were small, with a diameter of 70 µm or less (mean = 46 µm ± 19.0sd) and lacking in any surface ornamentation. These small, smooth spores varied with respect to melanization, with spores exhibiting three main colors: hyaline, orange, and black (Figure 1A, 2A). Spores contained intact lipid contents, indicating viability. Variation in spore traits was not explained by meteorological factors such as wind speed, temperature or precipitation. In particular, maximum wind speed was not a strong predictor of spore size and the largest AM fungal spore (400 µm) was collected in September when wind speeds are at an annual low.

**Figure 1.**
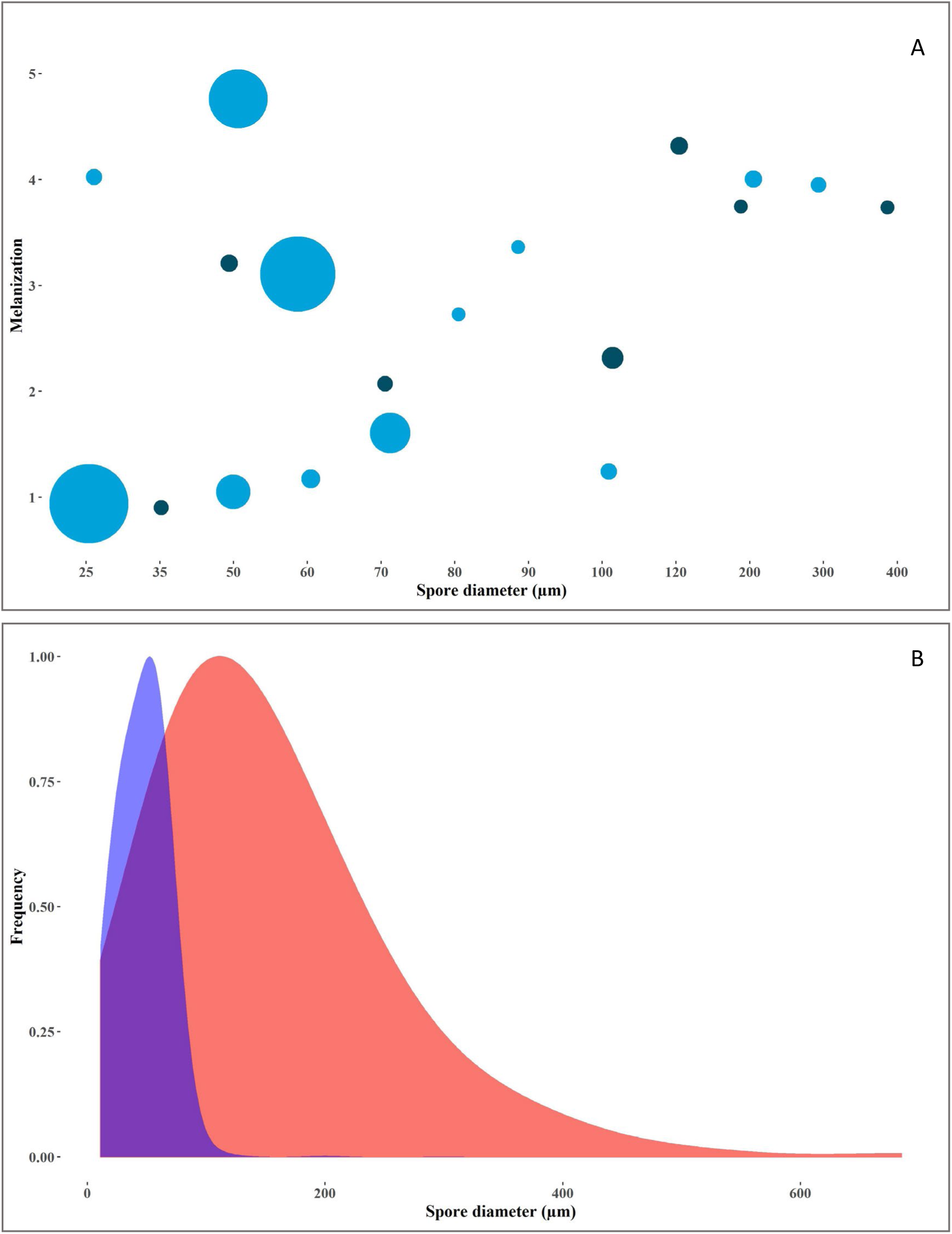
(A) Heatmap of traits of 47,162 aerial AM fungal spores collected over the course of one year. Size of point indicates the abundance of spores in a particular trait space. Dark blue = ornamented spore, light blue = no surface ornamentation. (B) Distribution of spore diameter for aerial AM fungal spores (blue) and average spore diameter for all described Glomeromycotina fungi (red) from data in Aguilar-Trigueros et al. (2019).

**Figure 2.**
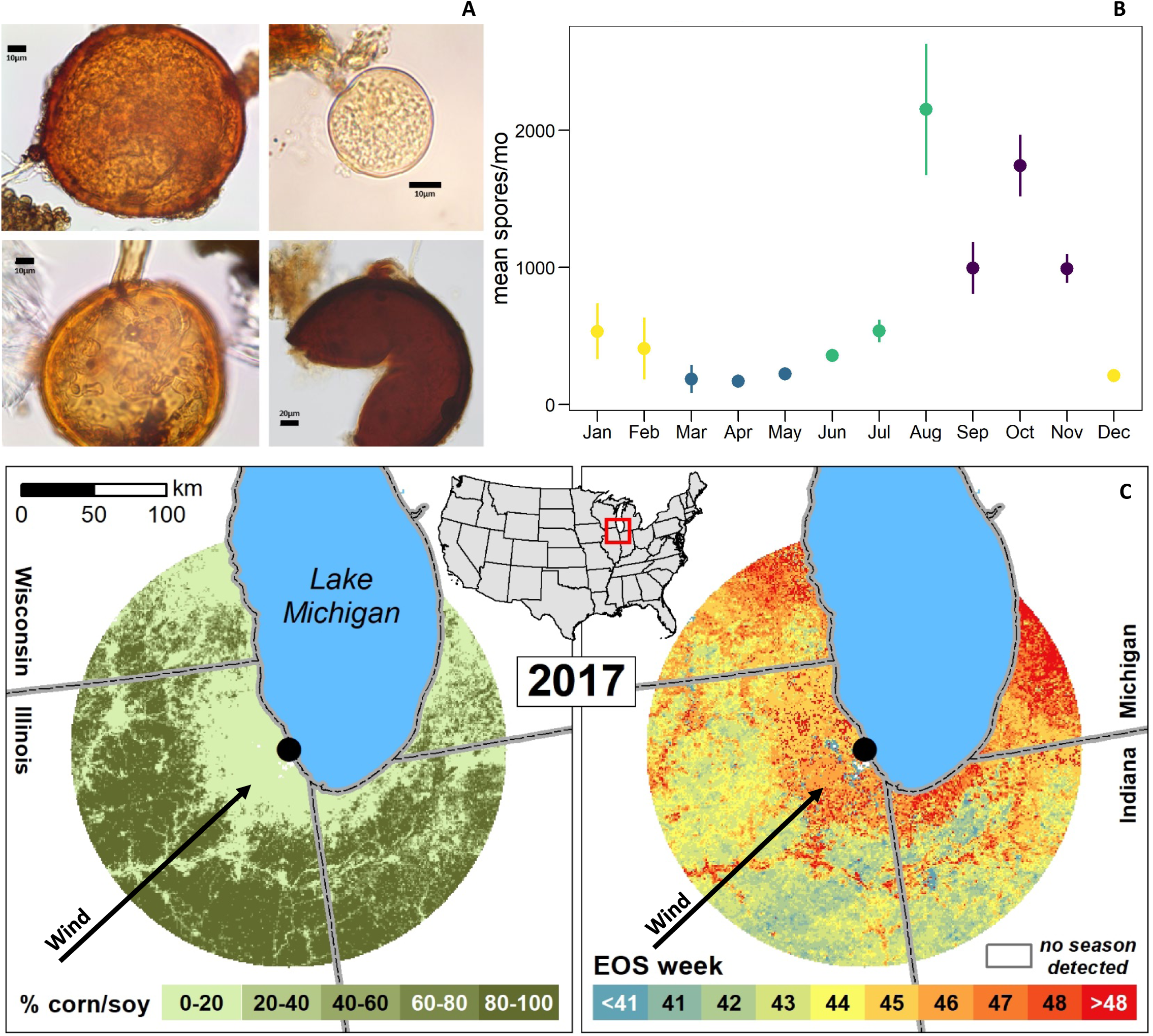
(A) Commonly encountered aerial AM fungal spores (clockwise from top left *Claroideoglomus etunicatum, Archaeospora trappei, Septoglomus constrictum*, and *Glomus microcarpum*), and (B) monthly mean ± standard error for spore abundance categorized by seasons (yellow = winter, blue = spring, green = summer, purple = fall). Lower-left panel in (C) shows corn or soy cover per 1 km^2^ pixel developed using USDA National Agricultural Statistics Service 2017 Cropland Data Layer (2019), while the lower-right panel illustrates the week during 2017 when dominant vegetation completed senescence (i.e. reached end-of-season, or EOS) for a 150 km buffer around study site.

Compared to all described Glomeromycotina fungal spores (Aguilar-Trigueros *et al.*, 2018), the size distribution of aerial AM fungal spores is narrower and centered on a smaller median diameter (Figure 1B). Spores of all described AM fungi range in size from 11 µm to 685 µm (mean = 153.5 µm ± 96.9sd); species forming spores smaller than 25 µm and larger than 400 µm were not present in our aerial samples. The overall relative distribution of aerial AM fungal spore diameter was significantly different from that of all Glomeromycotina spores (Kolmogorov-Smirnov: p < 0.0001). The overwhelming majority of aerial AM fungal spores in our dataset (99%) had an observed diameter of 70 µm or less. According to data from published species descriptions, 27% of AM fungal species form spores with minimum diameters of 70 µm or less (Aguilar-Trigueros *et al.*, 2018). These small-spore species belong to 17 of the 36 Glomeromycotina genera, including *Acaulospora, Ambispora, Archaeospora, Bulbospora, Claroideoglomus, Diversispora, Dominikia, Entrophospora, Funneliformis, Glomus, Pacispora, Palaeospora, Paraglomus, Redeckera, Rhizophagus, Sclerocystis*, and *Septoglomus*. Of these genera predicted to demonstrate the potential for aerial dispersal, 8 were detected in our aerial samples using morphological methods and 5 were detected using DNA-based techniques (results described below).

### Temporal patterns in dispersal

Spores of AM fungi were present in aerial samples during every month of the year, including winter (Figure 2B). A strong temporal pattern in AM fungal spore abundance was observed, with peak aerial dispersal of AM fungi occurring from August to November (Figure 2A). Minimum AM fungal spore capture occurred in March (50 spores/month) and the maximum occurred in August (3,661 spores/month). Interestingly, aerial AM fungal spore abundance was slightly negatively correlated with monthly maximum wind speed (Pearson r = - 0.592). Remote sensing data indicates that the primary windward vegetation in the 150 km radius region surrounding the sampling location is agricultural fields consisting of corn and soy and urban ecosystems (Figure 2C). Furthermore, remote sensing data indicate that, in 2017, windward vegetation reached peak senescence primarily prior to the 42^nd^ week of the year, or early October (Figure 2C).

In addition to temporal patterns in overall spore abundance, the six most dominant morphospecies also exhibited unique patterns with respect to when they were predominantly present in the air. In the year in which we sampled, *Archaeospora trappei* spores were primarily observed in August and *Glomus microcarpum* spores gradually increased to exhibit peak abundance in October (Figures 3A and 3B). *Glomus deserticola, Rhizophagus intraradices*, and *Diversispora spurca* spores had variable abundance over the course of the year, while *Acaulospora spinosa* was primarily observed in the air in July (Figures 3C-3F).

**Figure 3.**
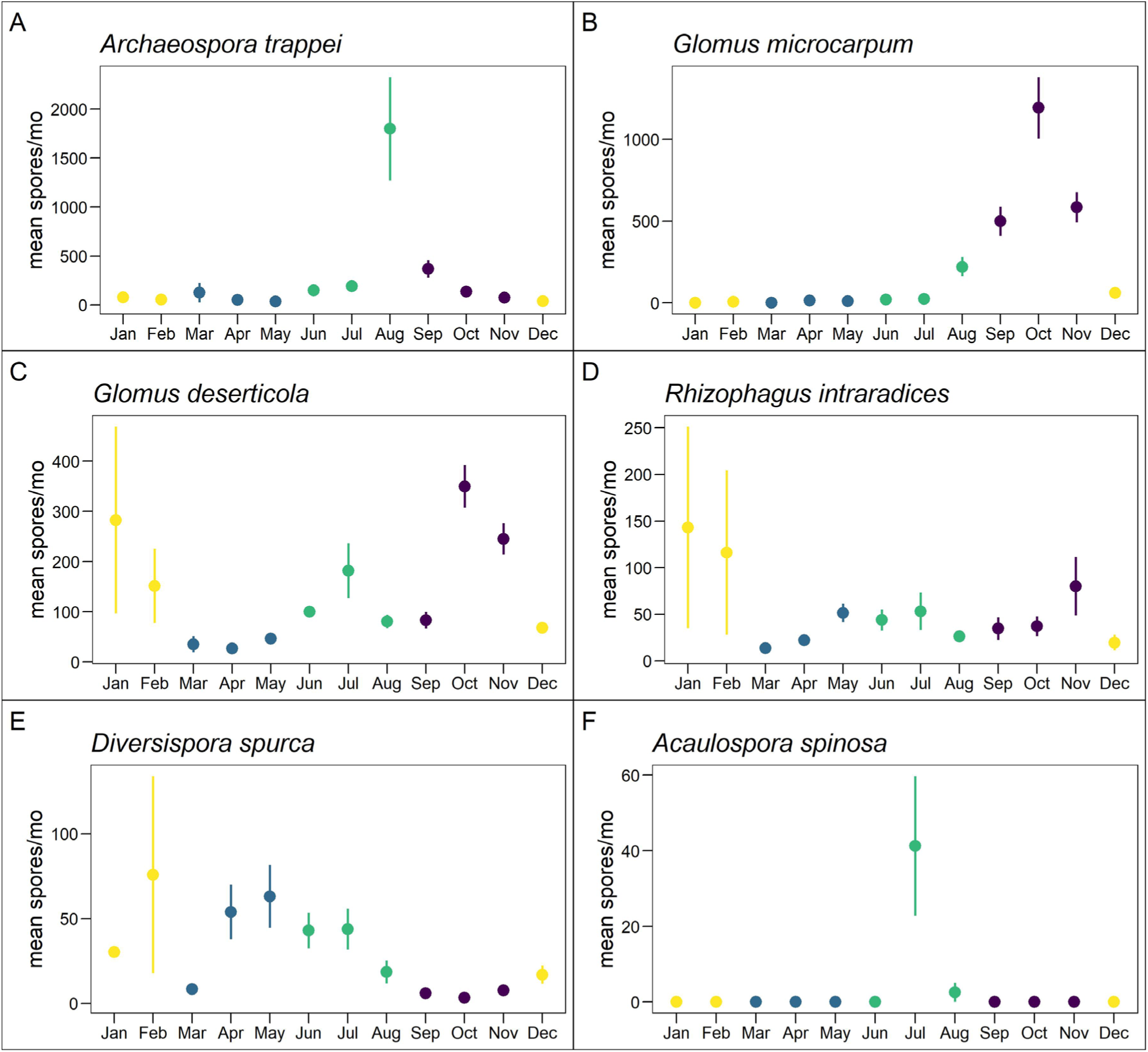
Temporal patterns in aerial spore abundance for the six most abundant AM fungal spore morphological species. Monthly mean ± standard error is presented for each species and further color coded by seasons (yellow = winter, blue = spring, green = summer, purple = fall).

### Spore species and virtual taxa in aerial samples

A total of 20 unique morphological species were observed over the course of one year with an average of 5 observed per month (min =4/month, max=8/month). Nine genera were represented including *Acaulospora, Ambispora, Archeospora, Claroideoglomus, Diversispora, Funneliformis, Glomus, Rhizophagus*, and *Septoglomus*, but *Glomus* species were the most common (Table 1; Figure S3). Species richness of aerial AM fungal spores did not differ significantly between months or seasons (data not shown), but spore community structure, accounting for both species identity and relative abundance of spores, shifted strongly between seasons (Figure 4A). Community structure of aerial AM fungi differed significantly by season (MRPP: p=0.001, A = 0.2498) and pairwise analyses showed statistical differences among all seasons (Table S1), with the most stark change occurring from fall to spring.

**Table 1.** Morphologically defined AM fungal spore species detected in aerial samples. Colors indicate different Glomeromycotinan families.

|  |  |
| --- | --- |
| *Acaulospora foveata* | *Funneliformis mosseae* |
| *Acaulospora morrowae* | *Glomus macrocarpum* |
| *Acaulospora sp. 1* | *Glomus microcarpum* |
| *Acaulospora spinosa* | *Glomus sp. 1* |
| *Ambispora jimgerdemannii* | *Glomus sp. 2* |
| *Archaeospora trappei* | *Glomus sp. 3* |
| *Claroideoglomus*<br>*etunicatum* | *Glomus sp. 4* |
| *Claroideoglomus*<br>*lamellosum* | *Rhizophagus clarus* |
| *Diversispora spurca* | *Rhizophagus intraradices* |
| *Funneliformis geosporum* | *Septoglomus deserticola* |

**Figure 4.**
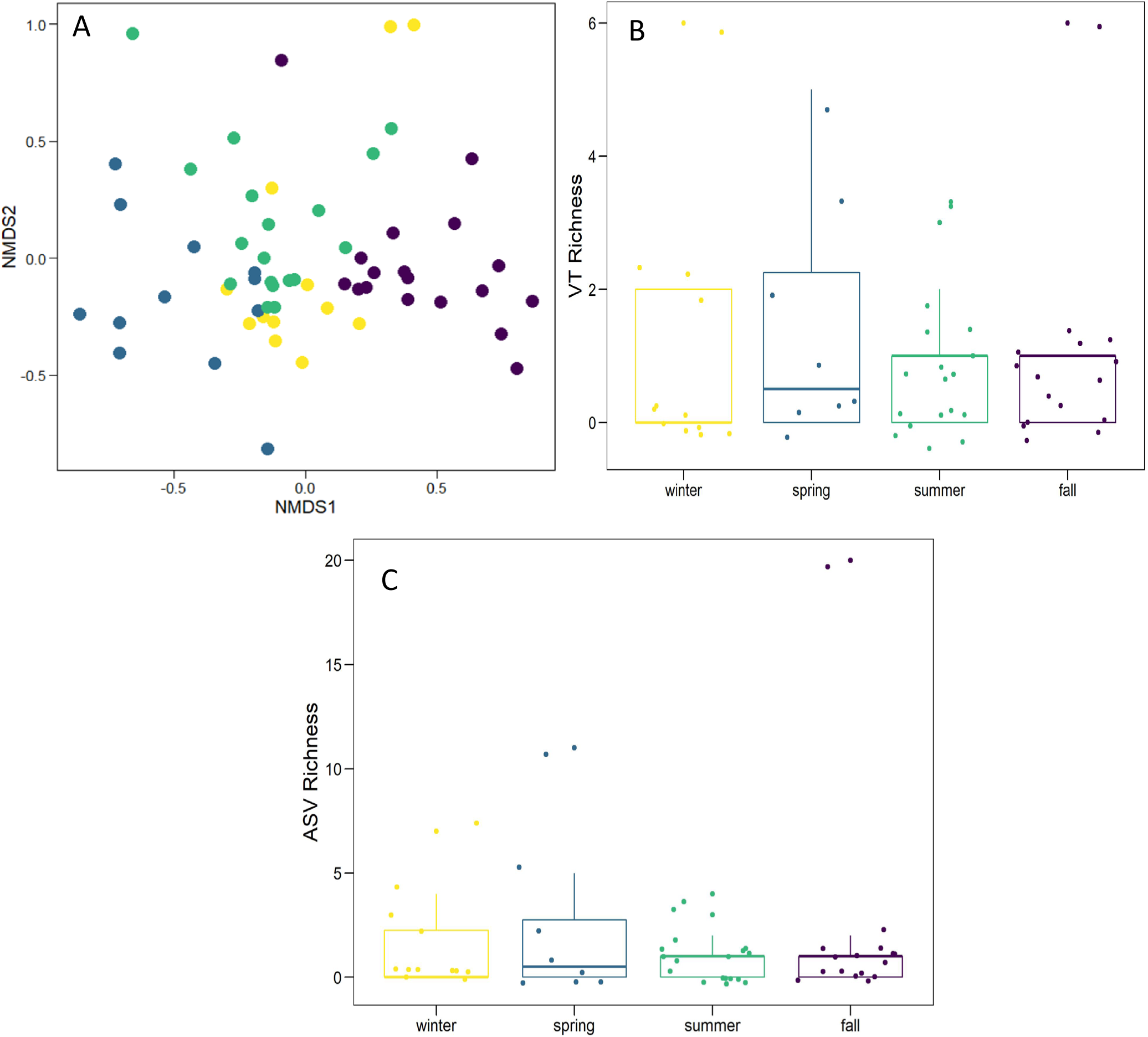
Temporal patterns in aerially dispersal of AM fungi as measured by (A) community structure of unique spore morphotypes and their relative abundances, (B) richness of virtual taxa (VT), and (C) richness of amplified sequence variants (ASVs). Yellow = winter, blue = spring, green = summer, purple = fall.

Over the course of one year, a total of 48 AM fungal ASVs were detected across all aerial samples. These ASVs matched 17 different VT belonging to five genera including *Claroideoglomus, Diversispora, Glomus, Paraglomus*, and *Scutellospora* (Table 2). Three genera that were observed using DNA-based methods - *Claroideoglomus, Diversispora* and *Glomus -* were also detected using spore morphology. Low amounts of AM fungal DNA in certain samples and sampling months led to variable amplification and sequencing depth (Figure S2), which precluded analyses of VT temporal patterns by month. However, pooling ASVs and VT by season allowed for qualitative presence or absence comparisons across sampling seasons (Figure 5).

**Table 2.**
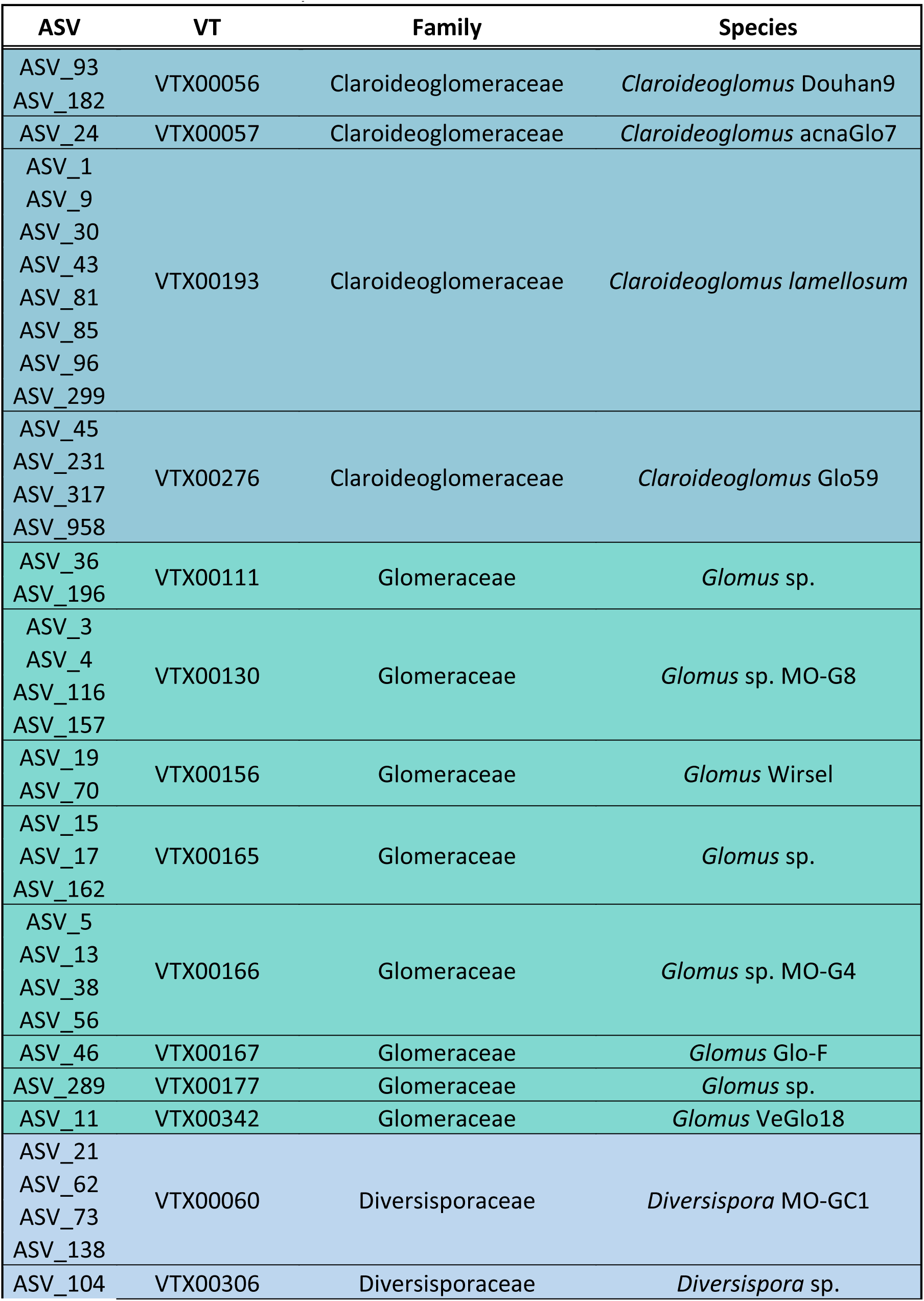
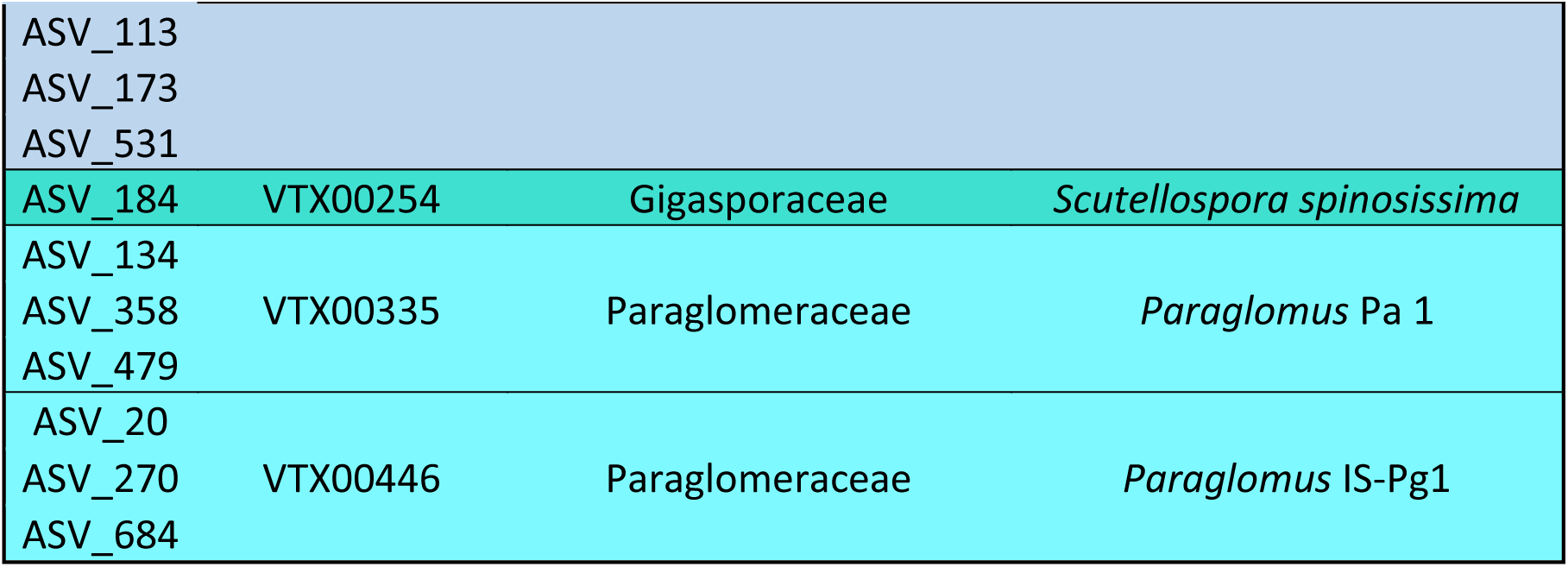
Virtual Taxa (VT) assignments from MaarjAM detected from whole dust. Colors indicate different Glomeromycotinan families.

**Figure 5.**
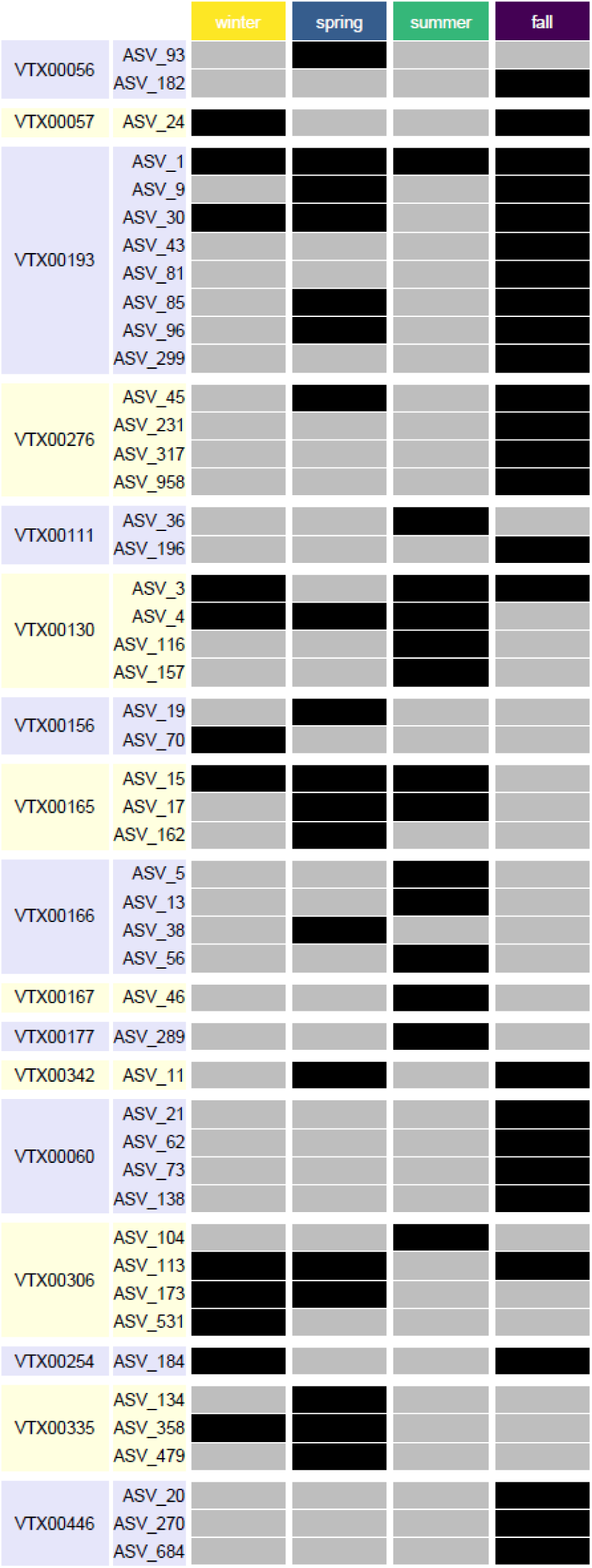
Table of AM fungal virtual taxa (VT) and amplicon sequence variants (ASV) detected in aerial samples across different season. Black boxes indicate present and grey boxes indicate absent.

The number of distinct VT was highest in the fall (11 VT), followed by spring (10 VT) and summer and winter (8 VT each) (Figure 4B). In the rarefied data however, summer exhibits the greatest diversity (8 VT), followed by fall (6 VT), winter (5 VT) and spring (4 VT). At the ASV level we observed the same trend with richness highest in fall (26 ASVs), followed by spring (19 ASVs), summer (14 ASVs) and winter (12 ASVs) (Figure 4C). In the rarefied data, fall shares the greatest ASV diversity with summer (14 ASVs each), followed by spring (7 ASVs) and winter (5 ASVs). Three VT, *Diversispora* VTX00306, *Glomus* VTX00130, and *Claroideoglomus* VTX00193 were present in aerial samples during all seasons. Four VT were only detected during a single season; *Diversispora* VTX00060, and *Paraglomus* VTX00446 were only observed in the fall, whereas *Glomus* VTX00167 and *Glomus* VTX00177 were only observed during summer. Several ASVs, representing genetic variation below the VT level, were detected in only a single season. Conversely, only ASV_1 which aligns to *Claroideoglomus* VTX00193 was present in aerial samples during all seasons.

## DISCUSSION

### Traits inform aerial dispersal

To our knowledge, this is the first study linking measured functional traits of AM fungi in the field with species-specific patterns of aerial dispersal. In support of H1, over the course of one year, we found a large amount of AM fungal spores present in the air at 20 m and these spores were more likely to exhibit traits, namely spore size, that facilitate aerial dispersal. Aerial AM fungi had smaller average spore size and range of size variation compared to the distribution of size for all described Glomeromycotinan species. Across all spores collected over the course of one year, spore diameter ranged from 25 µm to 400 µm, though larger spores were rare and most aerial spores were smaller than 70 µm. Trait-based predictions using spore size as a proxy for aerial dispersal capability at 20 m indicate that nearly 1/3 of described AM fungal species (representing ½ of all genera) fall within the spore size range for possible aerial dispersal. This indicates that the prevalence of aerial dispersal in AM fungi is perhaps greater than previously suspected and not limited to arid ecosystems (Egan *et al.*, 2014; Bueno & Moora, 2019). Furthermore, the diversity of AM fungal species with the capacity for long distance aerial dispersal is likely higher than previously thought (Warner *et al.*, 1987; Allen *et al.*, 1989; Kivlin *et al.*, 2014). These results do not necessarily contradict previous work suggesting low AM fungal abundance and diversity in air (Warner *et al.*, 1987; Egan *et al.*, 2014), but rather that longer sampling periods conducted during the appropriate season are required to illuminate patterns in aerial AM fungal dispersal. Because certain AM fungal species are more likely to disperse aerially than others based on differences in spore traits, these findings have implications for stochastic mechanisms that may drive AM fungal biogeographic patterns.

The relationship between fungal spore size and aerial dispersal capability has been studied for nearly a century in other fungi that form aboveground sporocarps (Buller, 1922; Norros *et al.*, 2014; Pringle *et al.*, 2015). Spore size as a predictive trait for AM fungal dispersal could be particularly informative because plasticity in spore size is comparatively large for the Glomeromycota and because spore size as a trait is phylogenetically conserved across broad AM fungal groups (Aguilar-Trigueros *et al.*, 2018). For example, early diverging AM fungi belonging to the Paraglomeraceae and Archaeosporaceae consistently form small spores, and both of these families were represented in our aerial collections. In certain AM fungal families such as Glomeraceae, Acaulosporaceae, and Ambisporaceae, spore size can vary up to three orders of magnitude, so species-level size information is required to make predictions regarding dispersal. It is possible that the predominance of small AM fungal spores present in aerial samples is the result of a predominance of small spore species present at the source of the spores. The source locations of spores for our 20 m aerial collections is not known and we did not assess concomitant soil communities. However, prior research reports that regional soils, depending on the level of anthropogenic disturbance, contain diverse communities of AM fungi (e.g. *Acaulospora spinosa, Entrophospora infrequens, Claroideoglomus claroideum, Claroideoglomus lamellosum, Funneliformis mosseae, Glomus fasciculatum, Racocetra fulgida, Cetraspora pellucida*) representing a broad spectrum of traits, including size, ornamentation, and color (Liberta & Anderson, 1986; Dhillion & Anderson, 1993; Middleton *et al.*, 2015; Koziol *et al.*, 2018).

Further research identifying variation in functional trait space and how it relates to ecological processes like dispersal will improve our understanding of mycorrhizal ecology (Zanne *et al.*, 2019). Unlike spore size, variation in melanization or spore ornamentation were not related to aerial AM fungal dispersal in our study and could not be compared broadly to all Glomeromycota due to a lack of available trait data for all described species. Other traits such as spore shape, wall thickness, sporulation season, or relative amount of spores produced per species could also be important predictors of aerial dispersal capabilities, but these data do not exist in an easily accessible format. Further *in situ* work combining AM fungal communities with observed traits, as well as improved AM fungal trait databases, are required to move toward greater predictive frameworks and the development of AM fungal community assembly “rules”.

### Temporal shifts in aerial dispersal

In support of H2, we observed strong temporal patterns in aerial AM fungi with peak spore abundance occurring from August to November and aerial community structure (i.e. spore species identity and relative abundance) shifting seasonally. Lowest spore abundance was observed during the spring months of March through May, but AM fungal spores were still present in the air during all seasons including winter. Interestingly, abundance and community temporal patterns were not correlated with meteorological variables (i.e. wind speed, temperature, soil moisture); in fact, spore abundance was negatively correlated with monthly maximum wind speed. Two potential mechanisms could explain why more AM fungal spores were observed Aug-Nov and why different aerial spore communities were present during different seasons. First, seasonal variation in sporulation among AM fungal species could contribute to observed temporal patterns in aerial spores (Gemma *et al.*, 1989; Pringle & Bever, 2002); highest aerial spore abundance occurred in August, which could be a high sporulation period for *Archeaspora trappei*, the species that dominated samples during this month. The composition of aerial AM fungal communities could be dependent on seasonal sporulation patterns of specific species present in soils of the surrounding region (Dhillion & Anderson, 1993). Seasonal variation in sporulation could also potentially explain why certain AM fungal spore species dominated air samples at different times during the year. Because temporal patterns in ecology and a changing climate result in hierarchically complex dynamics (Ryo *et al.*, 2019), we would not expect observed seasonal aerial spore patterns to necessarily hold over multiple years; this study does, however, demonstrate the potential for aerial dispersal of particular AM fungal species to shift temporally over the course of one year.

Another potential reason for the observed temporal spike in spore abundance is the timing of dominant regional anthropogenic soil disturbance, which could elucidate a possible mechanism for the liberation of AM fungi from rhizosphere soil to enable aerial dispersal. In the regions upwind from our sampling location, managed ecosystems are dominated by corn or soybean monocultures that are harvested during fall months. As such, soil disturbance associated with harvest and subsequent fall tillage operations were likely concentrated when most crops reached suitability of harvest. Harvest cuts, shreds, and spreading of dead plant material requires operating heavy machinery across fields and many producers in the Midwestern US continue to utilize a fall tillage operation following harvest, sometimes immediately. By the end of August 2017, in northeastern Illinois, 99% of soybean was setting pods; by the third week of October, 98% of soybean was dropping leaves (i.e. mature) and 91% of soybean was harvested by mid-November (USDA National Agricultural Statistics Service Cropland Data Layer, 2019). Maturation of corn in northeastern Illinois in 2017 began in August with 97% mature by mid-October and 83% of corn harvested by mid-November (USDA National Agricultural Statistics Service Cropland Data Layer, 2019). Temporal concentration of soil disturbance in this region as a result of harvest and tillage activities between August and November in 2017 could have contributed to a fall peak in aerial spore abundance. Prior work has demonstrated how agricultural soil disturbance through harvest and tillage activities is linked to increased Aeolian dust and higher levels of aerial spores (Goossens *et al.*, 2001; Lee *et al.*, 2006). Both seasonal variation in species-specific sporulation and anthropogenic soil disturbances that liberate belowground AM fungi could interact to produce temporal variation in aerial AM fungal dispersal.

Contrary to H3, aerial samples contained a relatively high amount of diversity (20 morphological species and 17 VT) as detected using both morphological and DNA-based techniques. The two species lists generated by spore identification and SSU rRNA amplicon sequencing had certain similarities, such as *Claroideoglomus lamellosum* and other species belonging to *Claroideoglomus, Diversispora*, and *Glomus*. There were also noteworthy differences; for example, while no Gigasporaceae spores were detected, sequencing revealed one Gigasporacea VT, namely Scutellospora spinossisima VTX00254. Observed differences between spore species and VT could be due to several reasons. First, in order to have enough DNA for amplicon sequencing, DNA extraction was conducted on whole dust and not spores, so AM fungal VT could be represented from DNA in hyphae or colonized root fragments as well as spores present in aerial samples. In fact, discrepancies between spore communities and soil communities as revealed by sequencing methods have been previously reported (e.g. Hempel et al., 2007). Dispersal via hyphal fragments could represent an alternative strategy employed by species that form large spores, and would lie outside of predictions made using a spore trait framework. Second, it is possible that, during DNA extraction, not all spores were lysed to release DNA for extraction or not all spores contained enough DNA for analysis, which would also explain the observed high-levels of unspecific amplification. While AM fungal sequences can be obtained from as little material as a single spore (Schwarzott & Schüßler, 2001), in these studies all DNA used for PCR (or at least a very high proportion) was from AM fungi. In our dust samples the amount of DNA obtained from spores, provided it could be extracted, might have been at a very low relative abundance compared to DNA from other organisms, resulting in unspecific amplifications. Finally, because spore and DNA analyses were conducted on subsamples of each observation, it is possible that, due to amounts of spores and DNA that are considerably lower than in soils, the different methods did not reach sampling saturation and captured different taxa (Figure S2).

### A model for AM fungal dispersal

The results of this and other studies contribute to the development of a conceptual model to organize hypotheses and identify knowledge gaps regarding AM fungal dispersal (Figure 6). Because soil disturbance is a widespread global phenomenon (Amundson et al., 2015), it may be a primary mode of AM fungal propagule liberation (Step 1) from rhizosphere soils for subsequent movement by a variety of vectors (Step 2). The dominant type of disturbance, likely agricultural in our study, will vary depending on region and could be the result of other large-scale activities such as grazing, recreation, mining, or urbanization. In our study, three of the VT that were present in aerial samples during all seasons (*Diversispora VTX00306, Glomus VTX00130, and Claroideoglomus VTX00193*) have been previously detected in studies from agriculturally disturbed soils (Moora *et al.*, 2014; Vályi *et al.*, 2015). Furthermore, because anthropogenically impacted soils generally contain fewer AM fungal species, liberation of AM fungal propagules primarily via human disturbance could lead to the disproportionate spread of “weedy” AM species that act as poor mutualists (Koziol *et al.*, 2018). This process could contribute to anthropogenic homogenization of AM fungal communities, or telecoupling, interconnection across geographic space formed as a result of human-environment interactions in an increasingly globalized world (Kapsar *et al.*, 2019). It is important to note that dispersal is contingent upon the successful colonization (Step 3) of deposited propagules, which likely depends on the type of propagule that is moving as well as other biotic and abiotic deterministic factors that influence the formation of mycorrhizas. To further our understanding of how dispersal influences AM fungal community assembly and biogeography, more research at all three steps of dispersal are needed.

**Figure 6.**
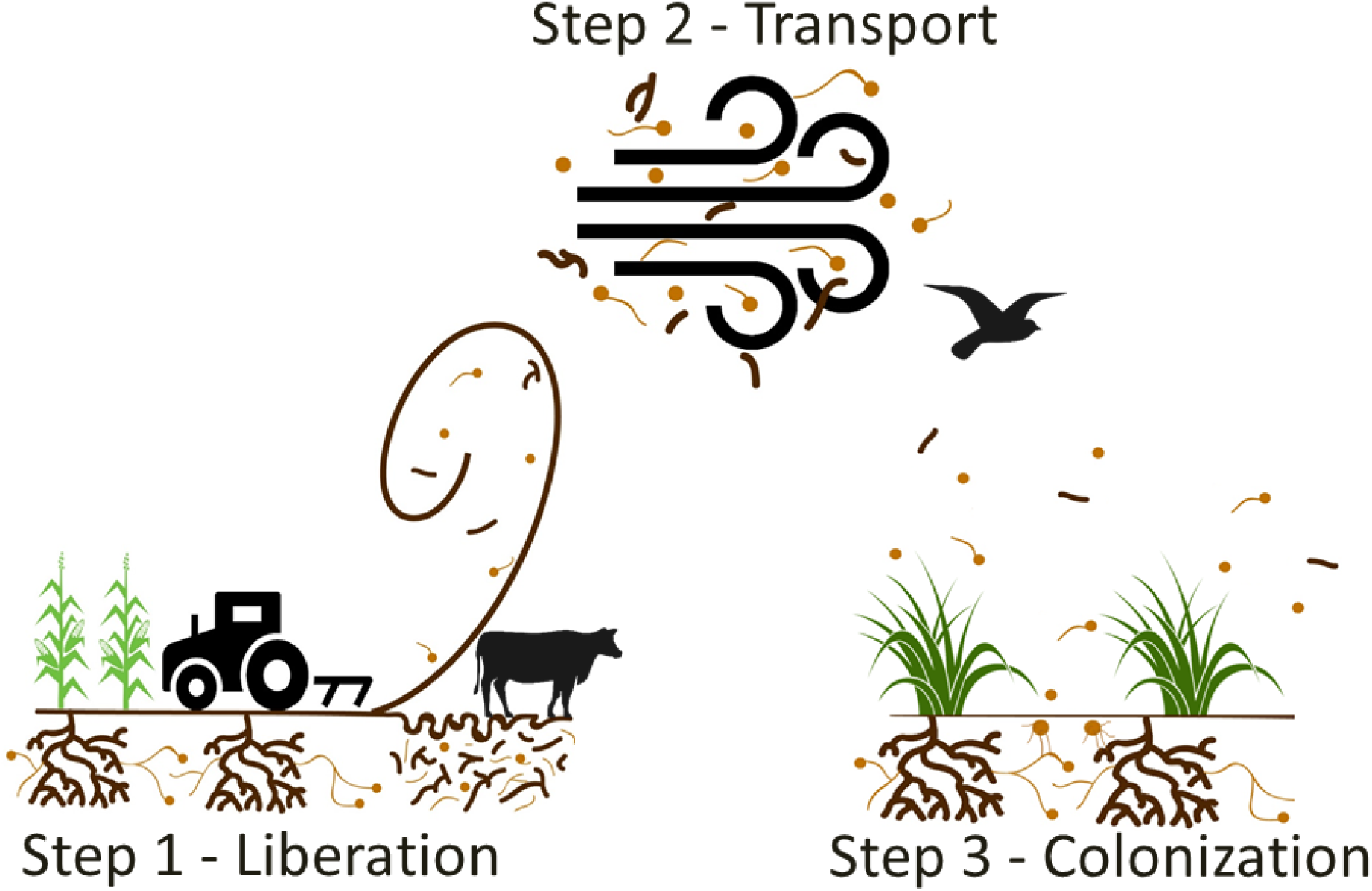
Conceptual model of hypothesized mechanisms of AM fungal aerial dispersal. Liberation, driven by anthropogenic soil disturbance, is required for mostly hypogeous AM fungal propagules to exit the rhizosphere. Transport of AM fungal propagules such as spores, hyphae, and colonized root fragments can occur via multiple vectors including wind, animals, or water. Successful dispersal is contingent upon deposition of AM fungal propagules into a new environment and subsequent colonization of host plant roots or an existing hyphal network.

This study provides evidence for a potential mechanism of the passive reestablishment and graduate increase in abundance of arbuscular mycorrhizas in a highly urbanized environment (Chaudhary *et al.*, 2019). Comparatively little research has been conducted in cities on arbuscular mycorrhizas, important symbionts for plants in urban ecosystems with the potential to enhance urban ecosystem services and sustainability (John *et al.*, 2016). Understanding how AM fungal communities reestablish after depletion from urban stressors could be key to conserving and restoring urban biodiversity in an effort to improve human well-being (Díaz *et al.*, 2019). Of interest to scientists who study mycorrhizal interactions, both in cities and beyond, is how to advise land managers and farmers regarding best practices in the management of AM communities and the potential benefits they confer to plant health. Recent commentary has broadened the discussion of *in situ* management of AM fungi (Rillig *et al.*, 2019), in part due to knowledge gaps regarding which species have the ability to passively disperse into sites without human intervention (Hart *et al.*, 2018). It is possible that, by identifying AM fungal species that are able to disperse aerially, native AM fungal inocula can be tailored to include species that cannot easily disperse. Investigating the mechanisms of AM fungal dispersal to improve current models of understanding will likely improve our ability to effectively manage mycorrhizas.

## ACKNOWLEDGMENTS

The authors are grateful to Ashlyn Royce and Kelly Velasquez for assistance with laboratory work. We thank Sarah Owens for consultation with next generation sequencing. Nancy Johnson, Thorunn Helgason, and three anonymous reviewers provided helpful comments on earlier drafts. VBC thanks Matthias Rillig for hosting an extended research leave that greatly facilitated the improvement of this manuscript. This work was supported financially by DePaul University and a National Science Foundation Grant (award DEB-1844531) to VBC.

## AUTHOR CONTRIBUTIONS

V.B.C conceived the study, collected data, conducted statistical analyses, and wrote the manuscript. S.N. conducted field sampling, processed laboratory samples, assisted with data collection, and provided editorial comments. C.E. assisted with next generation sequencing and provided editorial comments. M.A.S-H. performed the bioinformatical analyses and related analyses, contributed ideas, text, and editorial comments. J.K. developed the remote sensing-based phenology dataset and provided editorial comments.

