## Supplementary Material for "Trait-based aerial dispersal of arbuscular mycorrhizal fungi"

**SUPPLEMENTARY INFORMATION**

**Table S1.** Bonferroni corrected p-values for all pairwise comparisons of aerial AM fungal spore community structure among seasons.

| | **Fall** | **Spring** | **Summer** |
|---|---|---|---|
| **Spring** | 0.006 | | |
| **Summer** | 0.006 | 0.006 | |
| **Winter** | 0.006 | 0.024 | 0.006 |

**Figure S1.** BSNE passive dust samplers on rooftops for aerial AM fungal sampling. Photos were taken immediately after a winter and spring deployment (rain cover not shown).

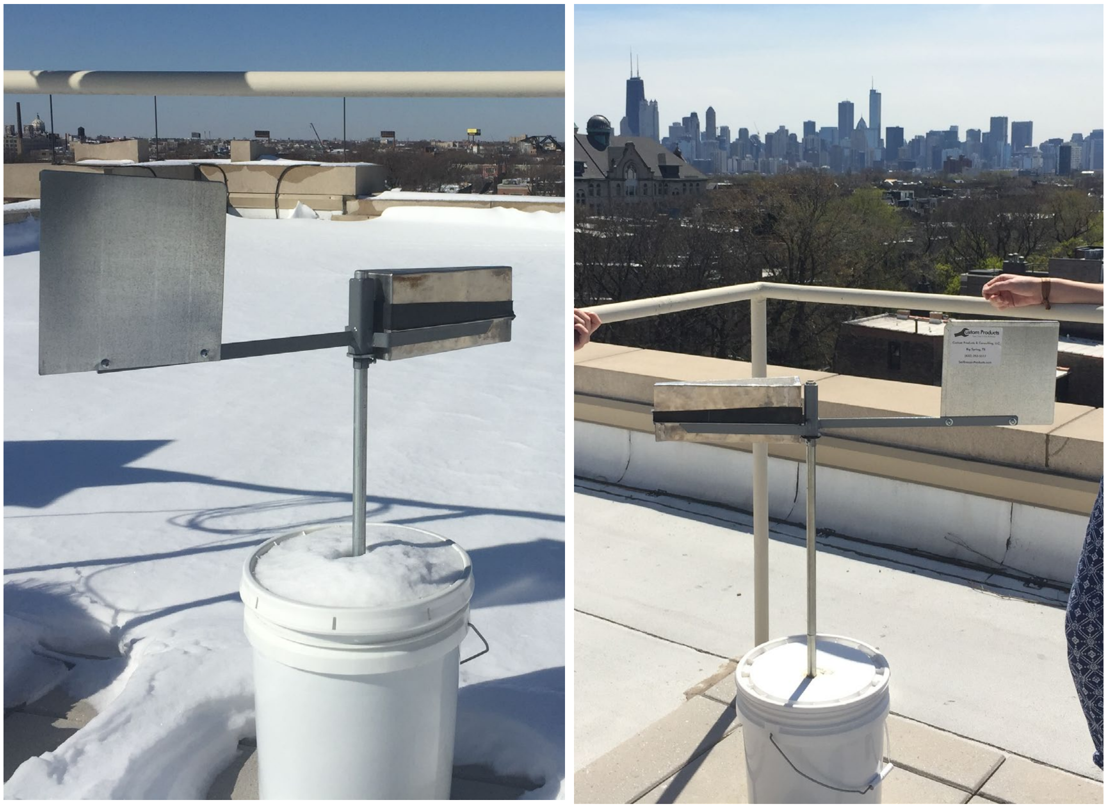

**Figure S2.** Species accumulation curves for aerial ASVs separated by season.

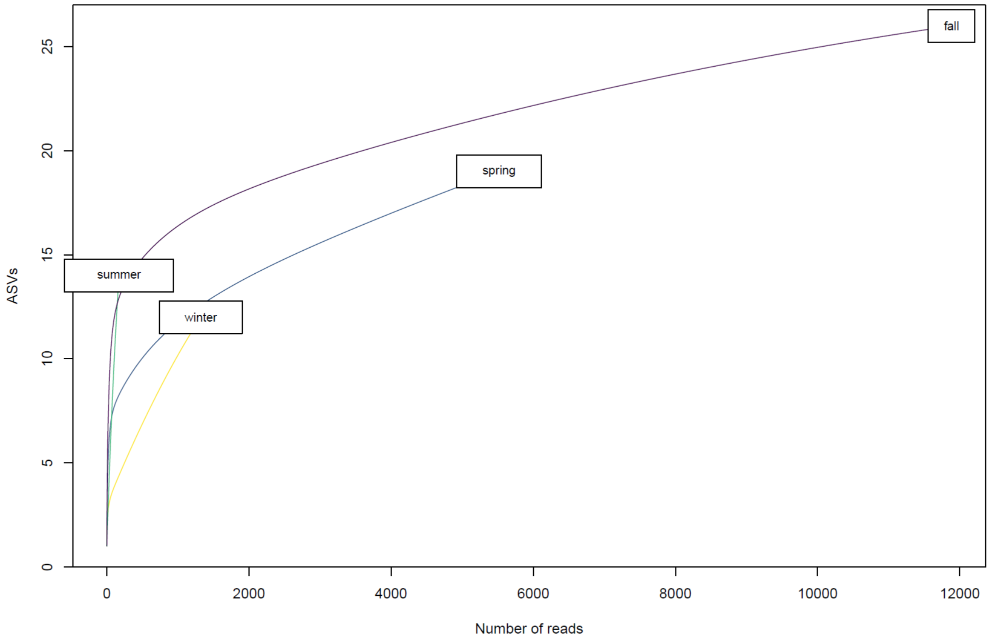


**Figure S3.** Species accumulation curves for morphological AM fungal spore species.

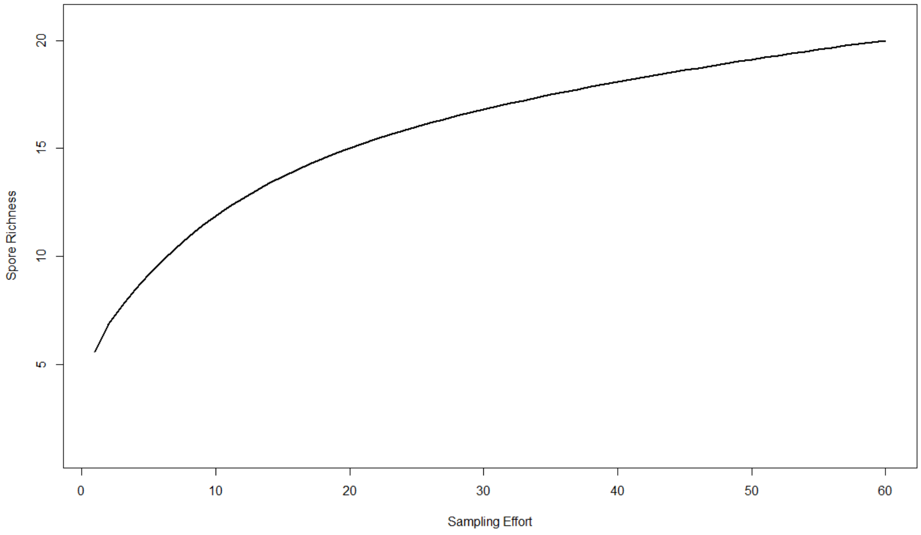
